# Effects of temperature and salinity on microbial degradation of bacterial necromass in urban river sediments in outdoor mesocosm experiments

**DOI:** 10.1101/2024.07.03.601918

**Authors:** Una Hadžiomerović, Daria Baikova, Iris Madge Pimentel, Dominik Buchner, Anna-Maria Vermiert, Philipp M. Rehsen, Verena S. Brauer, Rainer U. Meckenstock

**Affiliations:** Aquatic Microbiology, Environmental Microbiology and Biotechnology, Faculty of Chemistry, University of Duisburg-Essen, 45141 Essen, Germany; Centre for Water and Environmental Research (ZWU), Essen, Germany; Aquatic Ecosystem Research, Faculty of Biology, University of Duisburg-Essen, Essen, Germany; Department of Animal Ecology, Evolution and Biodiversity, Faculty of Biology and Biotechnology, Ruhr University Bochum, Bochum, Germany

**Keywords:** microbial self-recycling, microbial community, multiple stressors, stream sediments, food web

## Abstract

Microorganisms play a key role in the functioning of healthy river ecosystems because they consume carbon and nutrients from dead biomass derived from algae and other higher organisms, or “necromass”, to feed it back into the food web. This so-called microbial loop is known to be substantial in aquatic systems, but it is so far unknown to which extent microorganisms perform self-recycling, i.e. recycling of necromass derived from microorganisms, and how this process is affected by multiple stressors such as increased temperature and salinity. In this study, we investigated microbial self-recycling in the sediment of the urban river Boye within a food-web context before, during, and after a period of increased temperature and salinity using the outdoor flow-through mesocosm system “ExStream”. Rates of self-recycling were measured in additional microcosms with sediment and water from ExStream as an increase in ^13^CO_2_-concentration over time resulting from the microbial degradation of ^13^C-labelled microbial necromass, which was offered either in the form of intact but dead *Escherichia coli* cells, or lysed *E. coli* cells. The results showed that microbial self-recycling was highest in the first days of incubation suggesting that microbial necromass is an easily biodegradable carbon source. Increased salinity had no effect, but increased temperature or temperature and salinity strongly increased the rate of whole cell-necromass recycling compared to the unstressed control, while it had no effect on lysed cell-necromass recycling. After stressor removal, rates of self-recycling were not distinguishable from the unstressed control, showing the resilience of this community function. The compositions of the necromass-degrading communities were highly similar and little affected by stressor increase or release but changed during the course of the experiment. This indicated that both necromass types stimulated growth of the same organisms and that stressor levels were rather low for microorganisms and dominated by the effects of the seasonal variation in the Boye. The study suggests that microbial communities in urban rivers exposed to moderate levels of multiple anthropogenic stressors will be rather stressor-resistant and show fast recovery of community functioning.

## Introduction

Rivers are crucial for maintaining biodiversity, mitigating climate change effects and providing an array of ecosystem services such as water for drinking, agriculture, industry, energy and transport (Böck et al., 2018). The functioning of these ecosystems is to a considerable extent depending on processes in the sediments that are driven by microorganisms, which generally dominate aquatic ecosystems in terms of both biomass and abundance (Zinger et al., 2012). Microorganisms degrade necromass and other organic material and mineralise carbon and nutrients and are therefore crucial for the internal recycling of the contained elements within the riverine food web (Fenchel, 2008; Graham et al., 2014).

The importance of these microbial degradation processes for the internal recycling of elements and the functioning of aquatic food webs is well known and is referred to by the term “microbial loop”(Azam et al., 1983; Fenchel, 2008; Tranvik, 1992). In the ocean, the largest fraction of carbon and nutrients that are bound by primary production is not consumed directly by higher trophic levels, but it first becomes available to microorganisms as dead particulate or dissolved organic matter, i.e. necromass, to heterotrophic bacteria and archaea (Azam & Malfatti, 2007). These heterotrophic microorganisms are subsequently consumed by protists, and protists are consumed by higher trophic levels, which means that on the way from primary production to the top position of the food web, carbon and nutrients make a detour via the microbial loop (Hall & Meyer, 1998; Pomeroy, 2007). Yet, it was recently discovered that microorganisms do not only degrade algae-or plant- and animal-derived necromass but also necromass derived from other prokaryotic microorganisms (Bradley et al., 2018; Dong et al., 2018; Shoemaker et al., 2021). These studies showed that the extent of this microbial self-recycling may be considerable as well (Rillig et al., 2021). Moreover, a recent study on microbial self-recycling in drinking water discovered that microorganisms receiving microbial necromass reached higher cell concentrations than microorganisms receiving yeast extract, even though the amount of added organic carbon was lower with the addition of necromass than the yeast extract (Chatzigiannidou et al., 2018).

If the microbial self-recycling plays a major role in the functioning of microbial communities, it will also be important for maintaining river health. This is of major ecological importance, as rivers are increasingly threatened by anthropogenic activities (Reid et al., 2019) and global climate (Woodward et al., 2010). While quite some research has been performed in recent years to better understand the effects of multiple stressors on rivers (Dudgeon et al., 2006; Reid et al., 2019) and how rivers recover when stressors are released (Silva et al., 2024; Verdonschot et al., 2012), very little is known about the effects of multiple stressor increase and release on the structure and functioning of prokaryotic microbial communities in rivers (Romero et al., 2020; Shade et al., 2012), and in aquatic systems in general (Balian et al., 2007; Pedros-Alio, 2006; Zinger et al., 2012). In this study, we therefore aimed to elucidate the extent of microbial self-recycling for the functioning of microbial communities in rivers, and how microbial self-recycling differs in phases of river degradation and recovery. We based our investigation of the effects of multiple stressor increase and release on the Asymmetric Response Concept (ARC), which is a conceptual framework for a generic understanding of ecosystem degradation and recovery from multiple stressors (Vos et al., 2023). The ARC states that river communities during the degradation and recovery are governed by different assembly processes, i.e. the roles of the assembly processes during the two phases are asymmetric. During degradation, community assembly is mainly controlled by environmental filtering based on the different tolerances of species towards the acting stressors, while during recovery, community assembly is mainly controlled by dispersal and biotic interactions (Vos et al., 2023). As a consequence, the predictability of community composition during recovery is much lower than during degradation, because dispersal and biotic interactions are more subjected to stochasticity and may produce manifold community trajectories, whereas the outcome of environmental filtering is less variable. In accordance with the ARC, we hypothesise that the composition of prokaryotic communities at the end of a degradation phase is more predictable and thus less variable, whereas prokaryotic community composition at the end of a phase of recovery is less predictable and more variable. With respect to microbial self-recycling, we also hypothesize that recovery of prokaryotic community composition to its original state may not or only partly occur, but recovery of community function, i.e. microbial self-recycling, is more likely to occur because functions that were previously fulfilled by prokaryotic taxa that were lost during stressor application may be taken over during recovery by other, functionally redundant taxa (Louca et al., 2018; Wohl et al., 2004).

To investigate the extent of the microbial self-recycling and to test the hypotheses about multiple stressor increase and release, we used the outdoor experimental set-up “*ExStream*”, in which natural river water flows through multiple mesocosms filled with river sediment and larger river substrates, allowing for a replicated experimental design (Madge Pimentel et al., 2024; Piggott et al., 2015). We thereby focussed on the stressors temperature increase and salinization, as these are particularly threatening river ecosystem functioning worldwide (Berger et al., 2018; Canedo-Arguelles et al., 2013; Johnson et al., 2024). To test the effects of multiple stressors and release of stressors, river water flowing through the mesocosm was first left untreated to allow for acclimation of the communities. Then, it was either heated, salinized, or both, to cause degradation of communities, and then left untreated again to allow for recovery. Sediment samples were taken from the mesocosm at the end of each experimental phase to determine the composition of microbial communities. Additionally, sediment samples at the end of each experimental phase were incubated in microcosms to determine degradation rates of ^13^C-labelled bacterial necromass and to estimate the extent of the microbial self-recycling.

## Material and Methods

### Study site

The mesocosm experiments were conducted at the river Boye, an Emscher river tributary. The Emscher catchment is located in the Ruhr Metropolitan Region of Northern Germany. It was used as an open sewer until 1990s but underwent substantial restoration, so that no more sewage water has been discharged into the Boye stream network since 2017. Since 2021 it is officially considered as fully restored (EGLV, 2022; Winking et al., 2014, 2015)

### *ExStream* mesocosm experiments

*ExStream* is a highly replicated open-field mesocosm system (Madge Pimentel et al., 2024; Piggott et al., 2015) that was constructed next to the river Boye (51.5533°N, 6.9485°E). It consisted of 64 circular flow-through mesocosm channels of 3,5L each with an outflow in the middle. The mesocosms were filled with 1L of sediment from the river Boye mixed with 100 mL suspension of fine particulate organic matter from a small tributary stream, Spechtsbach, located close to the Boye (51.5627885 °N, 6.9150225 °E). The water from the Boye River was redirected to flow through the mesocosms. The water entering the system was divided into four spatial blocks that had two sedimentation tanks each to remove any larger particles from clogging the system. Each block was supplying water to 16 circular mesocosm channels what were set up randomly to avoid any block effects. For a more detailed description of the stup please see David et al. (2024); Mayombo et al. (2024). The experiment started on 4 March 2022. In the beginning of the experiment, all mesocosm received untreated water for 20 days from March 04-March 24 to allow for a phase of acclimation. Subsequently, a stressor phase was imposed for 14 days, from March 24 to April 07, where temperature and salinity were increased in a full-factorial design. Water temperature in the header tanks was increased by 4.5°C to achieve an increase of 3.5°C (mean ± SD: T_ambient_ = 8.7 ± 0.1 °C, n=4; T_heated_ = 12.1 ± 0.1 °C; n=2) in the mesocosms (see Supplementary table S1 and S2, Figure S1). Heating was interrupted twice due to equipment failures and high sediment loads, resulting in 9 days of effective warming. Salinity was increased by adding a NaCl-solution to the water supply, raising chloride concentrations by 154 mg/L above ambient levels (see Supplementary table S3, Figure S2). Salinity treatment was interrupted for 1.5 days due to heavy rainfall, resulting in 11 days of effective salinization. Finally, stressors were released in the recovery phase, lasting for 14 days from April 7 – April 21.

### Necromass production

#### Pre-cultivation

Bacterial necromass was produced from *Escherichia coli (DSM 13127*, DSMZ Braunschweig) cells as ^13^C-labelled necromass. To this aim, *E. coli* was cultivated on a nutrient agar (NA, Merck/Millipore) plate for 24 h at 37 °C, followed by plating onto a fresh agar plate and incubating for another 24 h at 37 °C. A single colony was then used to inoculate 100 mL of fresh M9 medium (Helmholtz Center Munich, see Supplementary Information S1) containing 0.4 % ^13^C-glucose (Cambridge Isotope Laboratories Inc, USA). The inoculated liquid culture was incubated in a 250mL baffled Erlenmeyer flask on shaker at 150 rpm at 37 °C for 24 h. Afterwards, it was used to inoculate a larger volume of 1900mL M9 medium, which was separated into five 1000mL baffled flasks (400mL in each) and incubated for an additional 24 h under the same conditions.

#### Washing

The 2000 mL liquid cultures were transferred to sterile 500 mL centrifuge containers and centrifuged at 3214 g for 10 minutes at room temperature (Eppendorf, model 5810 R, Wesseling-Berzdorf, Germany). The supernatant was discarded, and the pellet was resuspended in 100mL of sterile Milli-Q® water. The washing process of centrifugation and pellet resuspension was repeated three times in order to remove remnants of ^13^C-glucose from the medium. Washed labelled living *E. coli* cells were resuspended in 5 mL of Milli-Q® water.

#### Cell killing

The washed cell suspension was used to produce necromass both as dead but intact cells (“whole cell-necromass”) and as lysed cells (“lysed cell-necromass”). Whole-cell necromass was produced by killing *E. coli* cells with UV-light for 3h to simulate necromass resulting from intrinsic mortality. Two mL of the washed cells were diluted to a concentration of 10^6^ cells/mL and transferred to a sterile glass Petri dish containing a magnetic stir bar. The Petri dish was placed on a magnetic stirrer within a sterile bench (LabGard ES Class II, NuAire, Plymouth, USA) and exposed to UV-light (GE Lighting “Germicidal G30T8 30W”) for 3 h at 10cm distance from the light bulb, under constant stirring. Subsequently, cells were centrifuged, the supernatant was discarded, and the pellet was resuspended in 20 mL of Milli-Q® water. To verify that cells were killed, 0,05 mL of the re-suspension was streaked in duplicate on an agar plate and incubated overnight at 37°C after which, no growth was seen. The remaining whole cell-necromass was transferred into 1.5 mL tubes, flash-frozen with liquid nitrogen, and stored at -70°C until further use. Lysed-cell necromass was produced by lysing cells with a French Press to simulate necromass resulting from viral lysis. To this end, 3 mL of washed cells were subjected to 10 cycles of French press at 690 bar using the 20K Pressure Small Cell (Heinemann Labortechnik GmbH, Duderstadt, Germany). Approximately 33% of the volume was lost leaving approximately 2 mL of lysed cells. Because the French Press-treatment could not be performed sterile, lysed cells were diluted to 20 mL and exposed to UV-light for 1 h to kill all remaining life cell. 0,05 mL were plated on an NA agar plate and incubated overnight to rule out survival. The remaining lysed cell-necromass was flash-frozen with liquid nitrogen and stored at -70°C until use.

### Flow Cytometry

The production of whole cell-necromass using UV-light required a standardized densities of *E. coli* cells. Cell densities were determined in two technical replicates with a NovoCyte Flow Cytometer equipped with a NovoSampler Pro (Agilent, Waldbronn, Germany). Two times 200 µL of each sample were pipetted into a 96-well microtiter plate (Sarstedt, Nümbrecht, Germany). Samples were stained with 2 µL of SYBR Green I nucleic acid stain (Invitrogen, Darmstadt, Germany) diluted 1:100 in DMSO, and incubated for 13 minutes at 37°C in the dark. 50 µL of each sample were measured with a 488 nm argon laser at a flow rate of 14 µL min^−1^. volume. To prevent carry-over between samples, two blank samples of sterile filtered Milli-Q® water were measured before each sample at a flow rate of 66 µL min^−1^.

### Necromass degradation rates

Sediment samples were collected from *ExStream* mesocosms at the end of each experimental phase and transported in a thermobox to the lab on the same day. There, sediment was first sieved through a 1 mm-mesh to remove larger organisms and pebble stones. Then, 5 g of sieved sediment was added to a sterile 100 mL serum bottle and amended with 50 mL of Boye river water. Beforehand, this water was sterilized by filtration through a 0.2 µm-filter to avoid the introduction of organisms not originating from the sediment. Serum bottles were also amended with ^13^C-labelled necromass corresponding to a total amount of 2.5 × 10^9^ *E. coli* cells or 0,05mg of organic carbon. As the Boye water contained 5mg/l of dissolved organic carbon (DOC), the added necromass constituted about 20 % of the natural DOC-concentration. Serum bottles were sealed with a butyl stopper and amended with 500 µL of a 1M ^12^C-bicarbonate buffer, containing a natural atomic ^13^C fraction of 1.1%. The buffer was needed to increase the concentration of inorganic carbon and to allow for the determination of necromass degradation rates based on changes in the ^13^C/^12^C -ratio of CO_2_. Microcosms were then incubated for 60 days at the *in situ* temperature of the *ExStream* mesocosms. Necromass degradation rates were determined based on changes in ^13^C/^12^C -ratio of CO_2_ in the microcosms resulting from the degradation of the ^13^C-labelled necromass over time. To this, 2.5 mL of water was sampled from each microcosm for each timepoint with a 3 mL syringe equipped with a needle that was pierced through the stopper. A syringe filter with a pore size of 0.2 µm was installed between the syringe and needle to remove solids and the extracted volume was added in triplicates of 0.5 mL to sealed 12 mL Exetainer vials (Labco Limited, U.K.) that were flushed with dinitrogen gas and amended with 50 µL of 85% orthophosphoric acid. The Exetainer vials containing the samples were stored for up to 6 weeks until analysis with a G2131-i Isotope and Gas Concentration Analyzer (Picarro, USA) that measures δ^13^C in carbon dioxide. After each triplicate sample was measured, the first replicate was discarded and the other two δ^13^C values were averaged. The stable carbon isotope data were received as delta values (δ^13^) in ‰ and converted into an isotope-amount fraction (x(^13^C)) in percentage according to (Coplen, 2011):

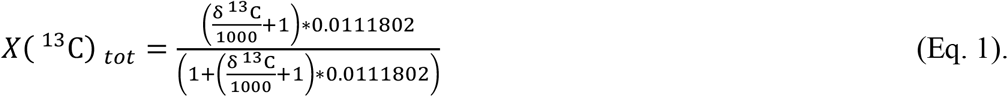

The isotope-amount fraction was then used to calculate the amount of ^13^C-bicarbonate in a sample, *n*_*2*_, using the following formula:

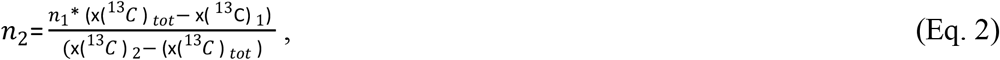

whereby *n*_*1*_ describes the amount of ^12^C-bicarbonate buffer in mM. x(^13^C)_tot_ describes the isotopic composition of the total bicarbonate in percent measured by Picarro, which is the sum of ^12^C-bicarbonate buffer and the bicarbonate already present in the sample. x(^13^C)_1_ describes the isotopic composition of ^13^C in the ^12^C-bicarbonate buffer in percent, and x(^13^C)_2_ describes the analyte isotopic composition of bicarbonate in percent, resulting e.g. from microbial activity. Next to the ^12^C-buffer and the ^13^C-necromass that was added to the microcosms, the microcosms also contained sediment and water, which contain DOC with natural abundance rate of ^12^C to ^13^C. Since this DOC can also be degraded by microbes, it will create predominantly ^12^CO_2_ that will influence the ^13^C/^12^C ratio that we measure to detect necromass degradation. Accordingly, in order to avoid underestimation of necromass degradation that is caused by degradation of natural DOC, we also measured DOC degradation rates in separate mesocosms without necromass addition to correct the calculations. In these microcosms without necromass, a bicarbonate buffer was added with a ^13^C-content of 10 atom percent (Schulte et al., 2019). To correct Equation 2 for the amount of ^12^C-bicarbonate produced from DOC-degradation, the corrected isotope amount fraction Equation 2 had to be adjusted by taking the DOC production into consideration.

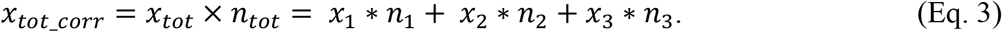

Here, *x*_*3*_ is the isotopic composition of ^12^C-bicarbonate in percent produced from DOC degradation and *n*_*3*_ is the produced amount of ^13^C-bicarbonate in mM. *n*_*tot*_ is the total amount bicarbonate in the sample. The values used for Equation 4 are provided in table 1. The corrected formula for the amount of ^13^C-bicarbonate then becomes

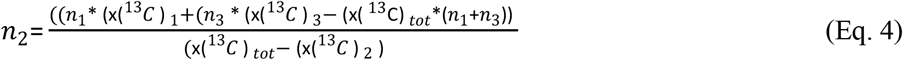

Rates of necromass degradation were calculated by dividing the amount of CO_2_ produced within the first five days by the length of the time period.

**Table 1:** Parameter values used for Equation 3.

| Parameter | Value | Unit |
| --- | --- | --- |
| $n_1$ | 10 | mM |
| $x_1$ | 1.10000514121476 | % |
| $x_2$ | 99 | % |
| $x_3$ | 1.10000514121476 | % |

### DNA extraction and sequencing

We analysed the composition of the microbial communities in the microcosms at the end of the incubation period. To this aim, microcosms were opened and sediment was transferred into 50mL falcon tubes. Sediment samples were centrifuged for 10min, and the supernatant was discarded. The sediment was transferred to eppis and frozen at -70 until DNA extraction. DNA was extracted in technical duplicates from 0.5 g of sediment using the following procedure. First, the samples were lysed by mixing with 0.1 and 0.5-mm diameter glass beads, 100 μL of Proteinase K, 5 μL of RNase A, and 900 μL of TNES buffer (see Buchner (2022d) for buffer and reagents). The samples were then bead-beaten for 2 minutes at 2400 rpm in a Mini-Bead-Beater 96 (Biospec Products, Bartlesville, USA) and subsequently incubated for 20 minutes at 56 °C. The lysates were divided into duplicates, and DNA was extracted following the spin column protocol using a vacuum manifold (Buchner, 2022a).

DNA extracts were cleaned up with carboxylated-magnetic beads and PEG-NaCl buffer where 40 μL of DNA and 80 μl of clean-up solution were mixed and incubated at room temperature for 5min at 900rpm (Buchner, 2022b). The plate was placed on magnet for 2min to pellet the beads. After discarding the supernatant by pipetting, 100 μL of wash buffer was added per sample while on the magnet, incubated for 30 seconds, and the supernatant was discarded. This washing step was repeated, followed by a 5-minute incubation to dry the pellet. Subsequently, 40 μL of elution buffer was added to each sample and incubated at room temperature for 5 minutes at 900 rpm. The plate was placed on the magnet for 2 minutes, after which 30 μL of eluted DNA was transferred to a new plate and stored at -20°C.

DNA amplification used a two-step PCR. The first PCR was conducted in a 10 μL reaction volume per sample using the Multiplex PCR Plus Kit (Qiagen) with primers 515f/806r (Apprill et al., 2015), and 1 μL of DNA input. Cycling conditions were 5 min of initial denaturation at 95 °C, 20 cycles of 30 s denaturation at 95 °C, 90 s of annealing at primer specific temperature followed by 30 s of elongation at 72 °C, and a final elongation for 10 min at 68 °C. Following the cleanup of the first PCR product (Buchner, 2022c), 2 μL of DNA was used for the second PCR. Cycling conditions were 5 min of initial denaturation at 95 °C, 25 cycles of 30 s denaturation at 95 °C, 90 s of annealing at 61°C followed by 30 s of elongation at 72 °C, and a final elongation for 10 min at 68 °C. DNA concentrations were normalized to 2 ng/μL using a bead-based normalization protocol (Buchner, 2022c), and samples were pooled together to one final library. The library was concentrated on silica spin columns (Buchner, 2022b) and eluted in a final volume of 100 μL. Library concentrations were first measured on Qubit Fluorumeter 2.0 using the Qubit™ 1X dsDNA HS Assay Kit (Thermo Fisher Scientific, Waltham, MA, USA) and then on the Fragment Analyzer using the High Sensitivity NGS Fragment Analysis Kit (Advanced Analytical Technologies, Inc., Ankeny, IA, USA) and then sent for paired-end sequencing on the Illumina NovaSeq (CeGat Gmbh, Tübingen).

### Sequence analysis

Raw sequences were processed using the R-packages DADA2 version 1.22.0 (Callahan et al., 2016) and subsequently analysed using phyloseq version 1.38.0 (McMurdie & Holmes, 2013) Raw sequence processing included removal of primers, trimming of reads at 240 bases, and standard sequencing error correction, denoising and chimera removal of the DADA2 workflow. Final abundance table contained amplicon sequence variants ASVs. Taxonomy was assigned to ASVs using the SILVA database version 138.1 (Quast et al., 2013). Further processing of relative abundance tables comprised removal of rare sequences occurring less than 10^−5^ times in total and removal of ASV assigned to chloroplast and mitochondrial DNA. Subsequently, the ASV composition of technical duplicates where checked by visual inspection and then merged by calculating mean relative abundances. The five most abundant ASVs found in negative controls were removed from all samples. Further processing and visualization of sequences were performed using the R packages phyloseq, ggplot2 (Wickham, 2016), ggpubr (Kassambara, 2023), tidyverse (Wickham et al., 2019), and vegan (Oksanen, 2009).

### Statistics

Statistical analysis were done using R packages lme4 and lmerTest for fitting mixed-effect models, and emmeans for the post-hoc model. The effect of the stressor and experimental phase on the necromass degradation rate was tested using a linear mixed effect model followed by pair-wise post hoc comparison with multivariate *t*-adjustment of *p*-values. Factors “Phase” and “Treatment during stressor phase” were modelled as fixed effects. Replicate samples we included using replicate identifier as a random factor. Degrees of freedom were calculated using the Satterthwaite method. Because separate mixed-effect models were used for each necromass type, resulting *p*-values were corrected using Benjamini-Hochberg adjustment. The effect of stressor, experimental phase, and necromass type on microbial community composition in microcosms was tested using a PERMANOVA including all three factors. Observed differences were considered statistically significant at a *p*-value ≤ 0.05. All sequence and data analysis were done using R version 4.1.2

## Results

### Necromass degradation

Degradation of ^13^C-labelled bacterial necromass was measured in closed microcosms as increase in the concentration of ^13^CO_2_. In all microcosm incubations, addition of ^13^C-labelled bacterial necromass resulted in a strong increase in the concentration of ^13^CO_2_, showing mineralization of the carbon contained in the lysed cell-necromass and in whole cell-necromass (Fig. 1a,b). The ^13^CO_2_ production was clearly due to the degradation activity of microorganisms rather than due to due chemical decay processes, as no ^13^CO_2_ was produced in the sterile controls (red lines in Fig. 1a,b). The increase in ^13^CO_2_-concentration was particularly high within the first five days but slowed down over time, indicating that most of the added necromass was turned over already in the very beginning, which indicates a high degree of bioavailability of both types of necromass. Thus, necromass degradation was substantial in all experimental phases of *ExStream*, in all treatments, and for both necromass types.

**Figure 1.**
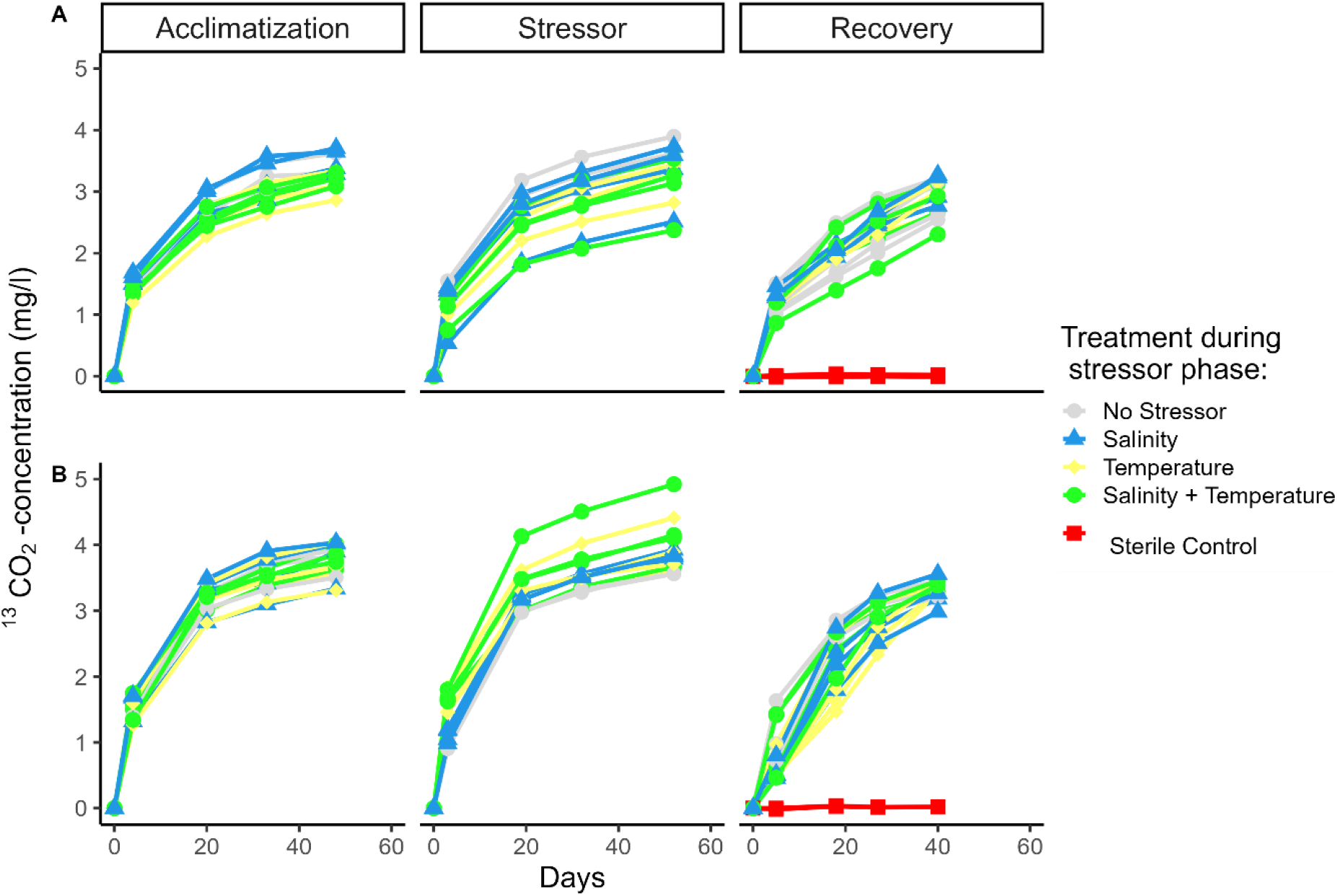
Changes in ^13^CO_2_-concentration over time in microcosms containing sediment and water from *ExStream*-mesocosms and amended with ^13^C-labelled lysed cell-(A) or whole cell-necromass (B). Symbols show means of two replicate microcosm incubations. Colours indicate stressor treatment applied during the stressor phase. Red squares show measurements in sterile controls.

Because the increase of ^13^CO_2_ in the microcosm incubations was generally highest in the beginning, maximum necromass degradation rates were calculated as the change in ^13^CO_2_-concentration over the first five days. The maximum degradation rates in the *ExStream* experiment were significantly affected by the experimental phase, the stressor treatment, and the necromass type (see Supplementary Table S4, S5, S6 and S7). At the end of the acclimation phase of *ExStream*, maximum degradation rates were highly similar for all mesocosms and for both necromass types, indicating that the experimental set-ups of the *ExStream*-mesocosms as well as of the microcosm incubations generate results of high reproducibility (Fig. 2a,b). At the end of the stressor phase, maximum degradation rates of lysed cell-necromass were not affected by any stressor treatment (Fig. 2a). Yet, maximum degradation rates of whole cell-necromass was significantly stimulated by the temperature increase and by the combined increase of temperature and salinity (Fig. 2b). At the end of the recovery phase, maximum degradation rates were again similar for all mesocosm and for both necromass types, but they were significantly lower compared to the previous experimental phases (Fig. 2a,b; Supplementary Table S4, S5, S6 and S7).

**Figure 2.**
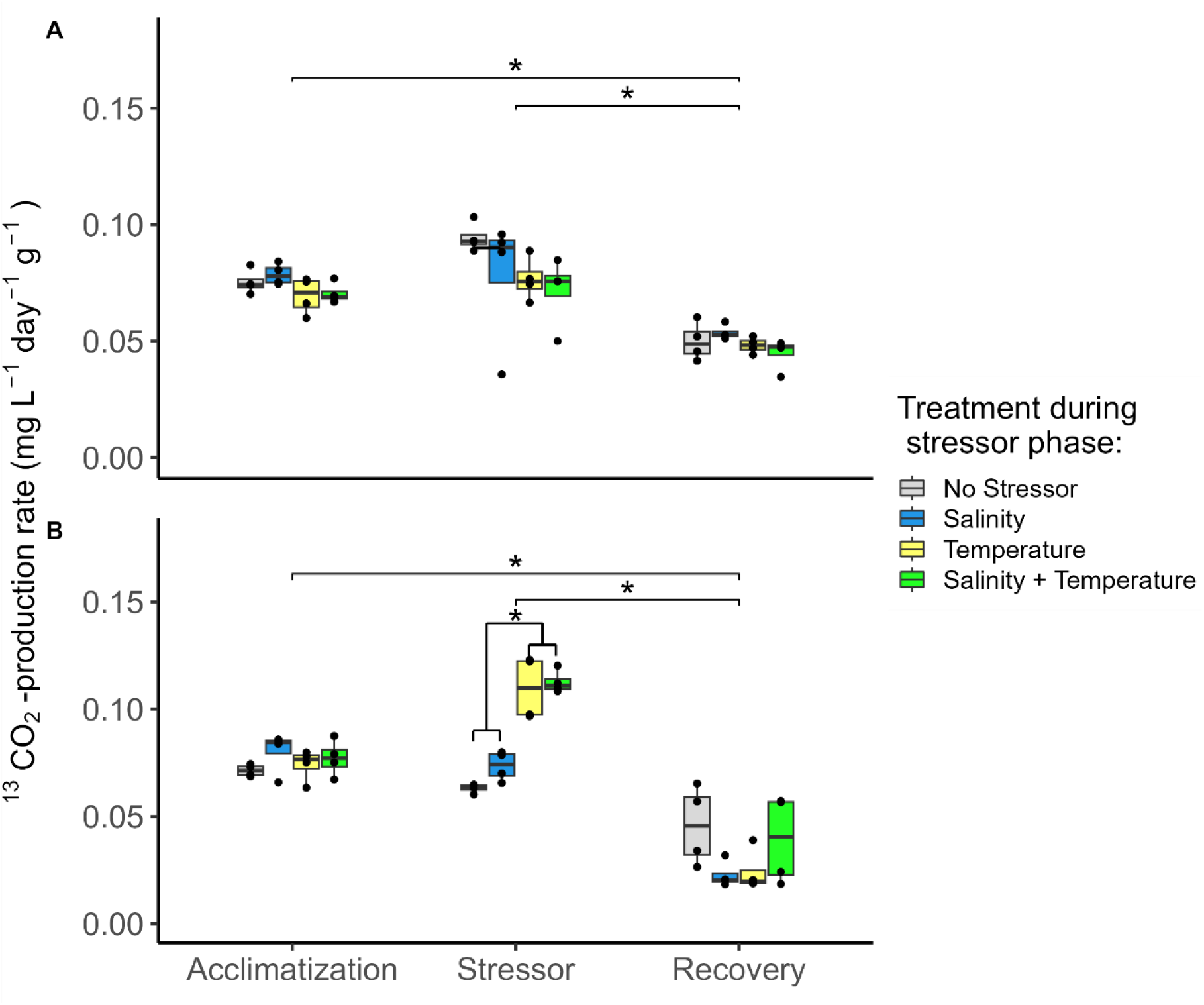
Rates of necromass degradation measured as ^13^CO_2_-production rate within the first five days in microcosms amended with lysed cell-(A) or whole cell-necromass (B). Boxplots represent median (horizontal line), interquantile range (boxes), and 1.5 x interquantile range (whiskers) of necromass degradation rates. Stressor treatment applied during the stressor phase is indicated by colour. Stars show significant differences in necromass degradation rate between experimental phases (A, B) and between mesocosms subjected to temperature or temperature and salinity increase and mesocosms subjected to salinity increase only or without stressor treatment (B; Linear mixed model followed by pair-wise post hoc comparison with multivariate *t-*adjustment of *p-*values; * indicates p<0.05).

### Microbial community composition

Microbial community composition was determined using 16S rRNA gene sequencing. To investigate the effects of multiple stressor increase and release on the necromass-degrading microbial communities, we determined microbial community composition at the end of the microcosm incubations containing lysed cell- and whole cell-necromass. These communities were compared with each other and to communities in additional microcosm incubations without necromass amendment in order to be able to relate observations to necromass rather than incubation effects. The results show that microbial community composition at the end of the microcosm incubations where significantly affected by the experimental phase in *ExStream*, stressor treatment, and necromass type, whereby the factor necromass type included lysed cell- and whole-cell necromass as well as no necromass (Fig. 3; Supplementary Table S8). In microcosm without necromass, communities appeared do differ between the experimental phases but not between stressor treatments (Fig. 3a). This indicates that the differences in these community compositions observed resulted from different community compositions in the mesocosm that were used as inoculum in each phase. In microcosms with lysed cell- and whole cell-necromass, communities appeared to differ particularly between stressor and recovery phase, and between the different stressor treatments applied in the stressor phase, but no difference was observable between the two types of necromass (Fig. 3b,c). Yet, necromass addition resulted in different community composition compared to no necromass addition in stressor and recovery phase (compare Fig. 3a and Fig. 3b,c).

**Figure 3.**
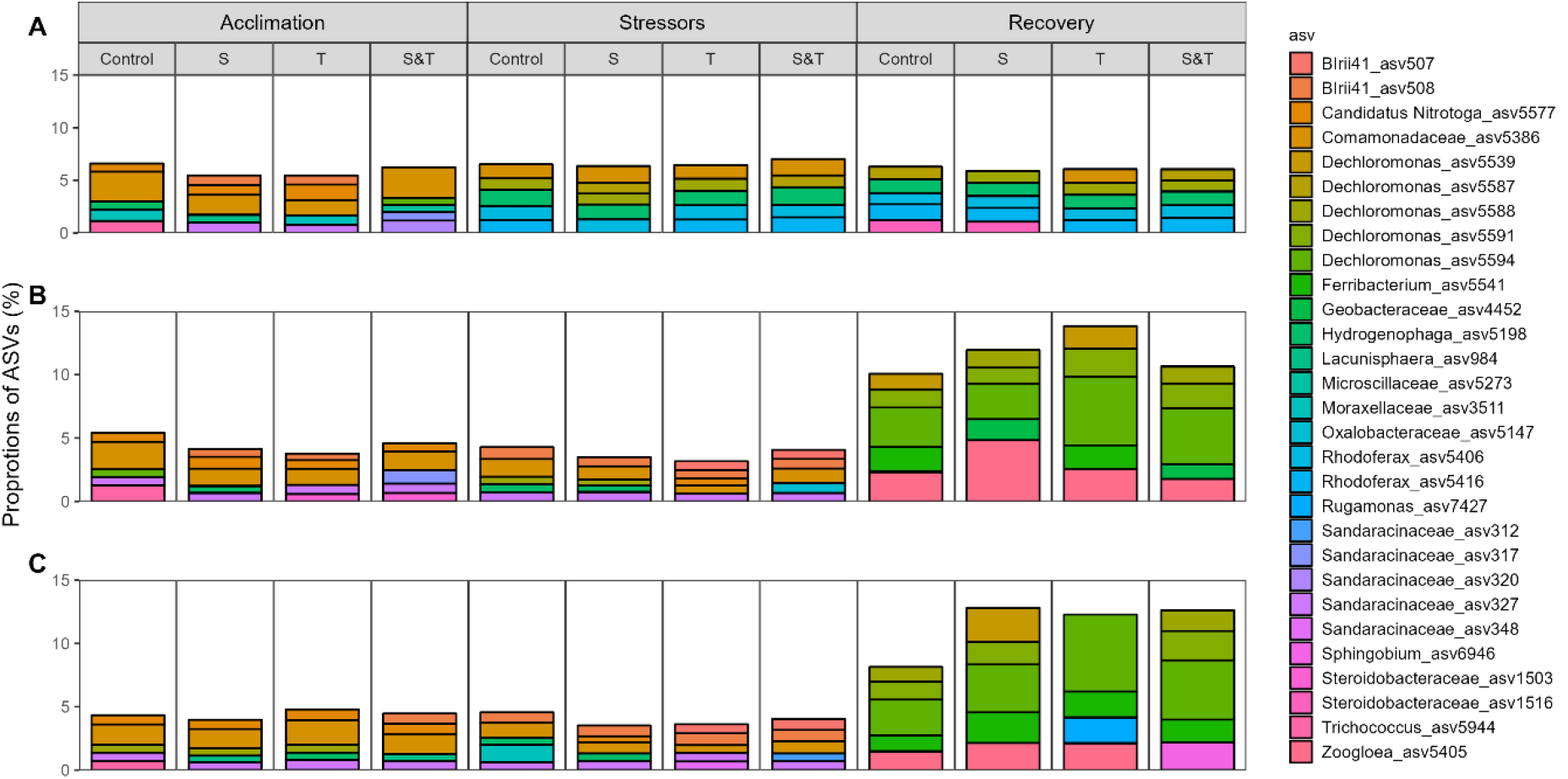
Proportions of the five most abundant amplicon sequence variations (ASVs) in each sample at the end of microcosm incubations without necromass (A), with lysed cell-(B), and whole cell-necromass (C). For visualization purposes, each bar shows the mean proportions resulting from four independent replicated *ExStream*-mesocosm experiments.

## Discussion

### Microbial self-recycling in river sediments

Our study indicates that microbial self-recycling in rivers may be substantial. Microbial self-recycling has recently been demonstrated in a number of studies where microbial communities live under extreme energy limitation, such as in marine sediments or in long-term experiments in closed systems (Bradley et al., 2018; Shoemaker et al., 2021). Microbial self-recycling has also been found in an anaerobic hydrocarbon degrading enrichment culture consisting of mainly two organisms, a hydrocarbon degrader and a necromass degrader. The necromass degrader was shown to degrade necromass amounts equalling 70% of the hydrocarbon degrader biomass, resulting in a high degree of hydrogen and nutrient recycling (Dong et al., 2018). Yet, self-recycling may also be substantial in other ecosystems where energy limitation is less extreme. In our study, the high necromass degradation rates observed at the beginning of the microcosm incubations show that bacterial necromass is a readily available carbon and nutrient source also for microbial communities in urban river sediments. The high bioavailability suggests that necromass is immediately turned over as soon as it becomes available, emphasizing the importance of microbial self-recycling for the cycling of elements in the food webs of such ecosystems. In order to be able to really estimate the proportion of self-recycling in overall microbial recycling in urban rivers, we would need to know the natural mortality of microorganisms, i.e. the rate at which microbial necromass becomes available. Yet, unravelling the extent of microbial self-recycling and the associated fluxes of carbon and nutrients is of major importance, because bacteria contain approximately 70 Gt of carbon and constitute about 15% of the global biomass (Bar-On et al., 2018). If self-recycling is substantial also in ecosystems that are not strongly energy-limited and that sustain a large food web with several trophic levels, this may mean that fluxes of carbon and nutrients through the food web are not clearly directional, and that a considerable fraction of carbon and nutrients is trapped in the self-recycling loop for a long-time.

### Degradation and recovery from temperature increase and salinization

The *ExStream* mesocosm experiment revealed only little effects of stressors on the composition and functioning of necromass-degrading communities. An increase in temperature and in temperature and salinity resulted in a significant increase in the degradation rate of whole cell-necromass, which is consistent with the general notion that temperature increases biological rates (e.g. (Arroyo et al., 2022; Rivkin & Legendre, 2001). No temperature effect was observable on the degradation rate of lysed cell-necromass. One possibility is that the higher degradation rate of whole cell-necromass with increased temperature indicates that this type of necromass is preferentially consumed by predatory bacteria or archaea and protists, which profit from higher temperatures as these increase chemical decay processes within the dead, but whole cells, making them more degradable. Yet, the composition of lysed cell- and whole cell-necromass degrading communities appeared to be very similar, suggesting that the observed differences may not be caused by different microbial taxa. Another possibility is that the lack of temperature effects on lysed cell-necromass degradation rate might not be real but due to a technical artefact. This might happen if increased temperature leads to a higher availability of ^12^C-organic material released from algae(Brauer et al., 2015), and therefore to a higher consumption rate of ^12^C-organic material. A higher consumption rate of ^12^C-organic material would not be detectable as increase in ^13^CO_2_-concentration, which was used in this study to measure necromass degradation rates. Thus, it is conceivable that increasing temperature in fact stimulated degradation rates of both necromass types, but that our method did not allow the detection of it.

Salinity increase did not lead to any changes in the degradation rate of lysed cell-or whole-cell necromass, nor did it affect community composition. This is consistent with another *ExStream* study that did not observe a salinity effect on microbial decomposition rates of cotton strips (Madge Pimentel et al., 2024). The effects of salinization on microbial diversity and process rates are generally not very clear, since both negative and positive effects have been reported(Gagnon et al., 2022; Kaushal et al., 2018; Martínez et al., 2020). In our study, it is possible that the amendment of 154 mg L^-1^ chloride ions in the *ExStream*-experiment was simply not high enough to cause salt stress in prokaryotes. The salinity stressor level was chosen as such that it would cause salt stress in macro-invertebrates and fish but at the same time, it would not be too detrimental for these organisms (Schröder et al., 2015). Another explanation is that microorganisms in the Boye are little susceptible to salinity stress because they are already adapted to higher levels. The Boye is a degraded river, which has been exposed to salinization resulting from coal mining in the past and which has only recently been restored (Madge Pimentel et al., 2024; Tröltzsch et al., 2020). Moreover, as an urban river, the Boye may also be more impacted by road salt application and treated waste water discharge that non-disturbed rivers. Indeed, a recent study comparing Boye with a non-degraded, near-natural river, showed that the communities of bacteria and other organism groups from the Boye showed much weaker responses to changes in salinity and several other environmental parameters than the communities from the non-degraded river Kinzig (Kaijser et al., 2024). The absence of the salinity effects on microorganisms in our study is thus consistent with other studies.

In contrast to stressor treatment, we observed strong differences in microbial community composition and a decrease in necromass degradation rate between the stressor and recovery phase of the *ExStream*-experiment. These differences were not caused by the release of the stressors, as they were also observed in the control mesocosm (see Fig. 2, 3 and Supplementary table S4, S5, S6 and S7), which means they were probably driven by seasonal changes happening in the Boye river. Indeed, strong changes within the river were observable in the the time period of these two phases, as water temperatures changed from 8.7°C to 10.9 °C. Moreover, we observed a strong increase in algal biomass covering the sediments in the mesocosms. The combination of algae and higher temperatures may have increased the availability of dissolved organic carbon, which then served as an easily degradable substrate for microorganisms, similar to the ^13^C-labelled necromass. As a consequence, microorganisms in the recovery phase may have degraded algal-derived ^12^C-DOC as well as ^13^C-labelled necromass, leading to an underestimation of the potentially possible necromass degradation rate is related to the measurement method.

### Asymmetric responses of community composition

In agreement with the Asymmetric Response Concept (ARC) we hypothesized, that community compositions of replicate incubations at the end of the stressor phase should be very similar because they are mainly controlled by the rather predictive process of environmental filtering. We also hypothesized that community compositions at the end of the recovery phase should be more dissimilar because they are more controlled by much less predictable processes of dispersal and biotic interactions (Vos et al., 2023). These hypotheses could not be confirmed in our study. Although communities were more generally more dissimilar in the recovery than in the stressor phase (see Fig. 3), this difference was also observed in the control incubations that had not been exposed to stressor increases, demonstrating again a strong influence of seasonal changes happening in the Boye river on the outcome of the *ExStream*-experiment, rather than influences related to stressor increase and release. One explanation might be again that the stressor intensities, which were chosen to be moderate in order to no be to detrimental for higher organisms, were not high enough to cause stress responses in microorganisms. In such a case, also no recovery responses from stress can be observed, meaning that data from microorganisms were insufficient to test the hypotheses derived from the ARC. Interestingly, however, along the three phases of *ExStream*, we observed a change from low community similarity at the of acclimation phase, to high similarity during the stressor phase, and then again to low similarity at the end of the recovery phase (compare sizes of red, blue and green clusters in Fig. 4). Using the ideas of the ARC, these changes may suggest that seasonal changes in the Boye during the “stressor phase” represented a much stronger stressor for the microbial communities than the stressors applied in the *ExStream*-experiment, masking potentially weak effects of the applied stressors. Likewise, the further seasonal changes during the “recovery phase” may have represented a much stronger stressor release effect than those applied in *ExStream*.

**Figure 4.**
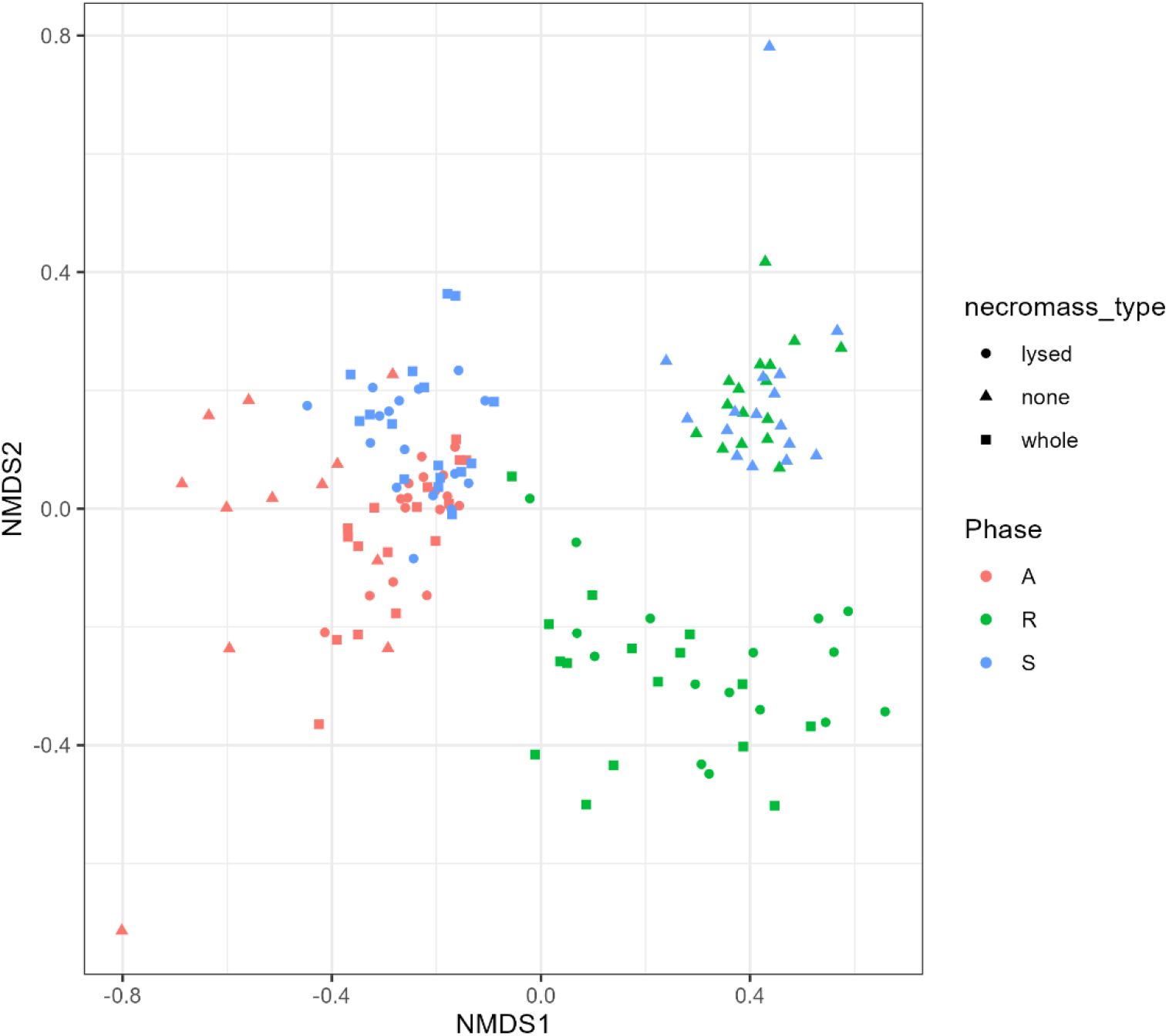
Non-metric multidimensional scaling of microbial communities sequenced at the end of closed microcosm incubations with no (triangles), lysed cell-(dots), and whole cell-necromass (squares) amendment. Microcosm contained sediment and water sampled from *ExStream*-mesocosms at the end of acclimation phase (“A”, red), stressor phase (“S”, blue), and recovery phase (“R”, green). Microbial communities cluster according to the experimental phases. Clusters size change from large during acclimation phase, to small during stressor phase, and back to large during recovery phase, indicating sequential changes in overall community similarity from low, to high, and back to low.

### Community composition versus community functioning

In a strict sense, we were not able to test our hypothesis that functions recover faster than community composition because stressor intensities in the *ExStream*-experiment were not high enough to cause microbial stress responses. However, we observed that community compositions changed along the course of the experiment, while community functioning in terms of necromass degradation was maintained, and always comparable to control mesocosms. Thus, we demonstrated a higher resistance of community functioning compared to community composition, and we observed that community functioning was resilient, as whole cell-necromass degradation rates in temperature-stressed mesocosm returned to the same level as control mesocosm after stressor release (Allison & Martiny, 2008). Resistance and resilience of microbial community functioning are usually attributed to the high functional redundancy of microorganisms, meaning that multiple taxa exist in a community that perform the same function. Thus, we can at least assume a certain degree of functional redundancy in the microbial communities of the Boye, which is the mechanism underlying the hypothesis of a faster recovery of functions compared to community composition.

Functional redundancy in microbial communities is well documented and wide (Louca, Jacques, et al., 2016; Louca, Parfrey, et al., 2016; Louca et al., 2018), and is probably rather high compared to higher organisms. For rivers exposed to moderate levels of multiple anthropogenic stressors, the results of this study therefore suggest that stressor release will result in fast recovery of microbial community functioning.

## Author contributions

**Una Hadžiomerović**: Data curation, Formal analysis, Investigation, Methodology, Project Administration, Software, Validation, Visualization, Writing – original draft, Writing – review & editing. **Daria Baikova**: Data curation, Investigation, Methodology, Project Administration, Validation, Writing – review & editing. **Iris Madge Pimentel:** Investigation, Methodology, Project Administration, Resources, Writing – review & editing. **Dominik Buchner**: Methodology, Project Administration, Resources, Writing – review & editing. **Anna-Maria Vermiert:** Investigation, Methodology, Project Administration, Resources, Writing – review & editing. **Philipp M. Rehsen:** Investigation, Methodology, Project Administration, Resources, Writing – review & editing. **Verena S. Brauer**: Conceptualization, Funding acquisition, Project administration, Resources, Supervision, Validation, Writing – original draft, Writing – review & editing. **Rainer U. Meckenstock**: Conceptualization, Funding acquisition, Project administration, Resources, Supervision, Validation, Writing – original draft, Writing – review & editing.

## Acknowledgements

The authors thank the *ExStream* 2022 team for running the mesocosm experiment, with a special thank you to student helpers Isabell Erdmann, Anna Mangels, Michaela Bojara, Julian Künkel and Jan Windhuis for all the help with the lab work, fieldwork and sampling.

This study was conducted within framework of the Collaborative Research Centre (CRC) 1439 RESIST (Multilevel Response to Stressor Increase and Decrease in Stream Ecosystems; www.sfb-resist.de) funded by the Deutsche Forschungsgemeinschaft (DFG, German Research Foundation; CRC 1439/1, project number: 426547801) and coordinated by B. Sures, D. Hering, and D. Grabner.

