## Supplementary Material for "Effects of temperature and salinity on microbial degradation of bacterial necromass in urban river sediments in outdoor mesocosm experiments"

Supplementary information S1

### M9 mineral medium

For 1 liter **M9 mineral medium** add to 867 ml sterile water:

| | | | |
|---|---|---|---|
| 100 ml | M9 salt solution (10X) | $Na_2HPO_4$<br>$KH_2PO_4$<br>NaCl<br>$NH_4Cl$ | 33.7 mM<br>22.0 mM<br>8.55 mM<br>9.35 mM |
| 20 ml | 20% glucose | glucose | 0.4 % |
| 1 ml | 1 M $MgSO_4$ | $MgSO_4$ | 1 mM |
| 0.3 ml | 1 M $CaCl_2$ | $CaCl_2$ | 0.3 mM |
| 1 ml | biotin (1 mg/ml) | biotin | 1 µg |
| ~~1 ml~~ | ~~thiamin (1 mg/ml)~~ | ~~thiamin~~ | 1 ~~µg~~ |
| 10 ml | trace elements solution (100X) | trace elements | 1X |

#### Stock solutions

| | | |
|---|---|---|
| **M9 salt solution (10X)** | $Na_2HPO_4$-$2H_2O$<br>$KH_2PO_4$<br>NaCl<br>$NH_4Cl$ | 75.2 g/L<br>30 g/L<br>5 g/L<br>5 g/L |

Dissolve the salts in 800 ml water and adjust the pH to 7.2 with NaOH. Add water to a final volume of 1 L and autoclave for 15 min at 121°C.

| | | |
|---|---|---|
| **20% Glucose** | 20% (w/v) glucose | 200 g/L |

For 500 ml stock solution add 100 g glucose to 440 ml water. ~~Autoclave for 15 min at 121°C.~~ Filter sterilize.

| | | |
|---|---|---|
| **1 M $MgSO_4$** | 1 M $MgSO_4$-$7H_2O$ | 24.65 g/100 ml |

For 100 ml stock solution dissolve 24.65 g $MgSO_4$-$7H_2O$ in 87 ml water. Autoclave for 15 min at 121°C.

| | | |
|---|---|---|
| **1 M $CaCl_2$** | 1 M $CaCl_2$-$2H_2O$ | 14.70 g/100 ml |

For 100 ml stock solution dissolve 14,70 g $CaCl_2$-$2H_2O$ in 94.5 ml water. Autoclave for 15 min at 121°C.

| | | |
|---|---|---|
| **Biotin (1 mg/ml)** | biotin (1mg/ml) | 50 mg/50 ml |

For 50 ml stock solution dissolve 50 mg biotin in 45 ml water. Add small aliquots of 1N NaOH until the biotin has dissolved. Add water to a final volume of 50 ml. Sterilize the solution over a 0.22-µM filter. Prepare 1 ml aliquots and store at -20°C.

| | | |
|---|---|---|
| **Thiamin (1 mg/ml)** | thiamin-HCl (1mg/ml) | 50 mg/50 ml |

For 50 ml stock solution dissolve 50 mg thiamin-HCl in 45 ml water. Add water to a final volume of 50 ml. Sterilize the solution over a 0.22-µm filter. Prepare 1 ml aliquots and store at -20°C.

| | | | |
|---|---|---|---|
| **100X trace elements solution** | EDTA | 5 g /L | 13.4 mM |
| | $FeCl_3$-6$H_2O$ | 0.83 g/L | 3.1 mM |
| | $ZnCl_2$ | 84 mg/L | 0.62 mM |
| | $CuCl_2$-2$H_2O$ | 13 mg/L | 76 µM |
| | $CoCl_2$-2$H_2O$ | 10 mg/L | 42 µM |
| | $H_3BO_3$ | 10 mg/L | 162 µM |
| | $MnCl_2$-4$H_2O$ | 1.6 mg/L | 8.1 µM |

Dissolve 5 g EDTA in 800 ml water and adjust the pH to 7.5 with NaOH. Then add theother components in the quantities mentioned below and add water to a final volume of 1
L. Sterilize the solution over a 0.22-µm filter.

| | | |
|---|---|---|
| 498 mg | $FeCl_3$ (anhydrous) | |
| 84 mg | $ZnCl_2$ | |
| 765 µl | 0.1 M $CuCl_2$-2$H_2O$ | 1.70 g/100 ml |
| 210 µl | 0.2 M $CoCl_2$-6$H_2O$ | 4.76 g/100 ml |
| 1.6 ml | 0.1 M $H_3BO_3$ | 0.62 g/100 ml |
| 8.1 µl | 1 M $MnCl_2$-4$H_2O$ | 19.8 g/100 ml |

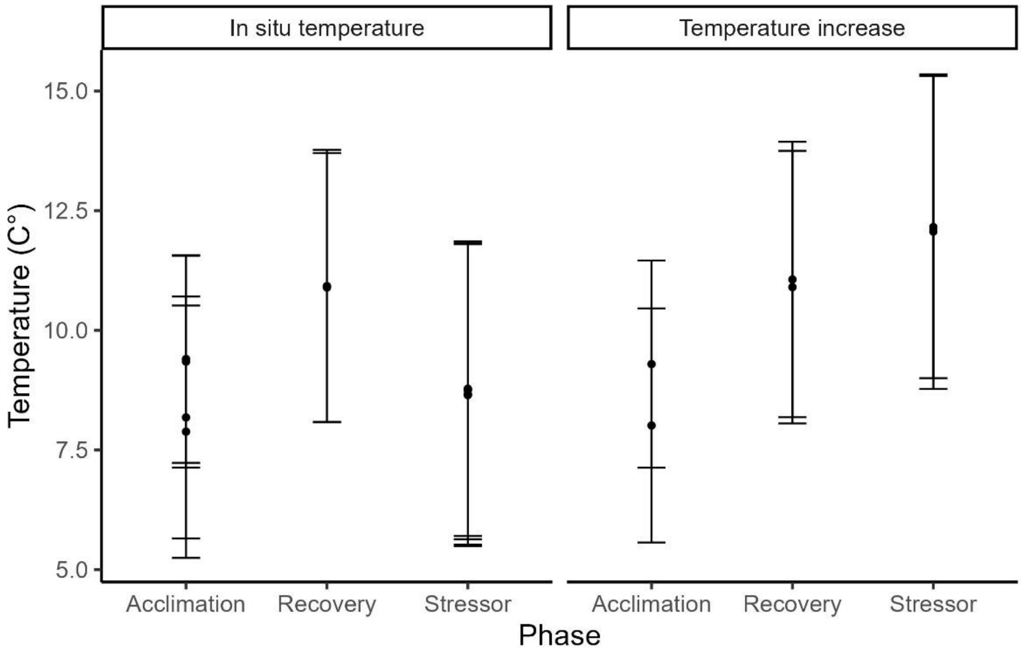


Figure S1. Temperature variation in channels during each phase of ExStream. Data are presented as mean temperature with standard deviation as error bars (note: only some channels had temperature loggers). In situ temperature mesocosms had 4 replicates with loggers while they were present in only 2 mesocosms with temperature increase.

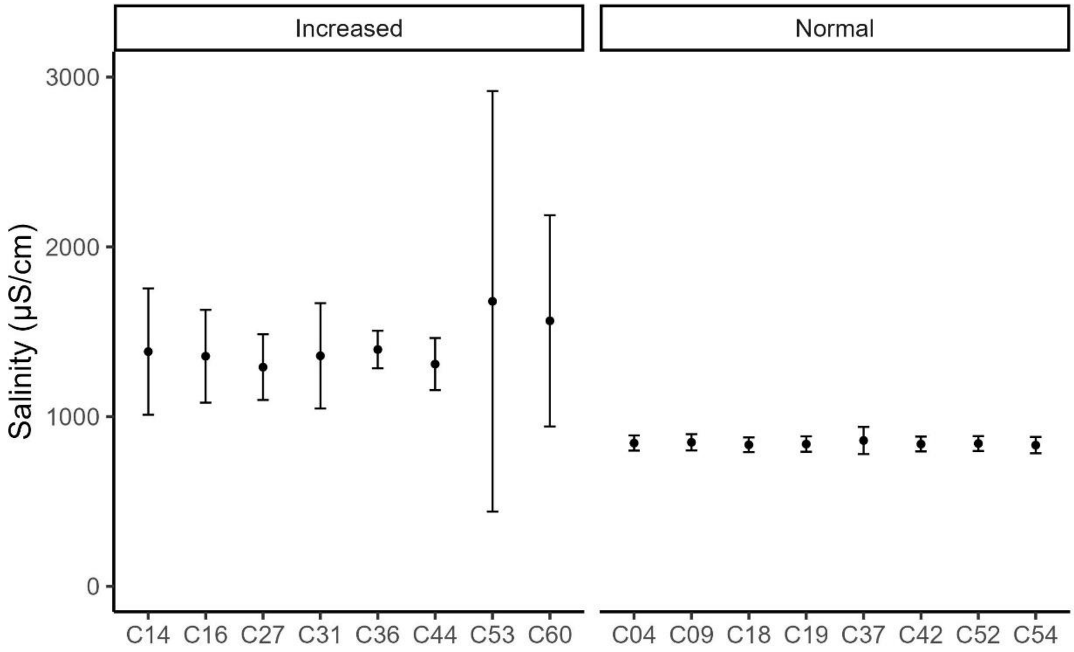


Figure S2. Salinity variation in 16 mesocosms(channels) during stressor phase used for microcosm creation. Data are presented as mean salinity with standard deviation as error bars.

**Table S1.** Summary statistics for temperature variation in channels of each phase of ExStream that had no temperature increase during stressor phase (*in situ*).

| Channel | Treatment | Phase | Mean temp | Sd temp | Max temp | Min temp | N |
|---|---|---|---|---|---|---|---|
| C25 | in situ temperature | Acclimation | 9.35 | 2.21 | 14.07 | 5.02 | 2276 |
| C29 | in situ temperature | Acclimation | 8.18 | 2.53 | 15.74 | 2.70 | 5510 |
| C35 | in situ temperature | Acclimation | 9.40 | 2.17 | 15.74 | 5.19 | 2276 |
| C41 | in situ temperature | Acclimation | 7.88 | 2.64 | 13.94 | 0.68 | 5510 |
| C29 | in situ temperature | Recovery | 10.93 | 2.84 | 19.09 | 3.99 | 4068 |
| C41 | in situ temperature | Recovery | 10.90 | 2.81 | 17.03 | 5.15 | 4068 |
| C25 | in situ temperature | Stressor | 8.71 | 2.65 | 15.40 | 3.60 | 3916 |
| C29 | in situ temperature | Stressor | 8.76 | 2.67 | 15.74 | 3.30 | 3916 |
| C35 | in situ temperature | Stressor | 8.82 | 2.61 | 15.40 | 2.57 | 3916 |
| C41 | in situ temperature | Stressor | 8.68 | 2.67 | 15.61 | 3.52 | 3916 |
| C25 | in situ temperature | Stressor, effective | 8.67 | 3.14 | 15.40 | 3.60 | 2308 |
| C29 | in situ temperature | Stressor, effective | 8.75 | 3.11 | 15.74 | 3.69 | 2308 |
| C35 | in situ temperature | Stressor, effective | 8.78 | 3.08 | 15.40 | 3.82 | 2308 |
| C41 | in situ temperature | Stressor, effective | 8.65 | 3.15 | 15.61 | 3.52 | 2308 |

**Table S2.** Summary statistics for temperature variation in channels of each phase of ExStream that had temperature increase during stressor phase.

| Channel | Treatment | Phase | Mean (°C) | Sd (°C) | Max (°C) | Min (°C) | N |
|---|---|---|---|---|---|---|---|
| C09 | Temperature increase | Acclimation | 9.30 | 2.16 | 14.88 | 5.06 | 2276 |
| C09 | Temperature increase | Recovery | 11.06 | 2.88 | 17.54 | 5.66 | 4069 |
| C09 | Temperature increase | Stressor | 11.12 | 2.95 | 19.47 | 5.06 | 3916 |
| C60 | Temperature increase | Acclimation | 8.01 | 2.45 | 13.85 | 2.40 | 5510 |
| C60 | Temperature increase | Recovery | 10.90 | 2.85 | 17.76 | 4.46 | 4068 |
| C60 | Temperature increase | Stressor | 10.82 | 3.18 | 19.09 | 3.09 | 3916 |
| C09 | Temperature increase | Stressor, effective | 12.16 | 3.16 | 19.47 | 7.03 | 2308 |
| C60 | Temperature increase | Stressor, effective | 12.07 | 3.29 | 19.09 | 3.09 | 2308 |

**Table S3.** Summary statistics for salinity variation in mesocosms of ExStream stressor phase that were used for creating microcosms.

| Microcosm | Chanell | Temp | Salt | Mean µS/cm | Sd µS/cm | Med µS/cm | Max µS/cm | Min µS/cm | N |
|---|---|---|---|---|---|---|---|---|---|
| 1 | C04 | normal | normal | 844 | 44 | 860 | 872 | 722 | 11 |
| 2 | C09 | normal | normal | 849 | 48 | 868 | 891 | 720 | 11 |
| 10 | C37 | normal | normal | 859 | 80 | 861 | 1062 | 721 | 11 |
| 11 | C42 | normal | normal | 838 | 44 | 856 | 870 | 722 | 13 |
| 5 | C18 | increased | normal | 834 | 43 | 849 | 868 | 720 | 11 |
| 6 | C19 | increased | normal | 838 | 45 | 858 | 871 | 720 | 11 |
| 13 | C52 | increased | normal | 842 | 44 | 860 | 870 | 722 | 11 |
| 15 | C54 | increased | normal | 832 | 48 | 856 | 871 | 720 | 13 |
| 3 | C14 | normal | increased | 1383 | 373 | 1315 | 2570 | 1033 | 13 |
| 4 | C16 | normal | increased | 1356 | 274 | 1323.5 | 2150 | 1072 | 12 |
| 9 | C36 | normal | increased | 1395 | 111 | 1426 | 1530 | 1174 | 13 |
| 12 | C44 | normal | increased | 1309 | 154 | 1318 | 1625 | 1063 | 11 |
| 7 | C27 | increased | increased | 1291 | 194 | 1260 | 1900 | 1096 | 13 |
| 8 | C31 | increased | increased | 1358 | 310 | 1307 | 2320 | 867 | 14 |
| 14 | C53 | increased | increased | 1679 | 1239 | 1329.5 | 5600 | 1143 | 12 |
| 16 | C60 | increased | increased | 1564 | 623 | 1404 | 3600 | 1265 | 13 |

**Table S4.** Model output for estimating the stressor and experimental phase effect on the necromass degradation rate of incubations with lysed cells.

| Parameter | Sum Sq | Mean Sq | NumDF | DenDF | F value | P-value |
|---|---|---|---|---|---|---|
| Treatment during stressor phase | 0.02 | 0.01 | 3 | 12 | 2.43 | 0.116 |
| Phase | 0.21 | 0.11 | 2 | 30 | 37.16 | < 0.001 |

**Table S5.** Pairwise comparison of experimental phase and stressor treatment of incubations with lysed cells.

| Phase1 | Treatment during stressor phase1 | Phase2 | Treatment during stressor phase2 | Estimate | SE | Df | t-ratio | P-value |
|---|---|---|---|---|---|---|---|---|
| A | SALT | S | SALT | -0.03 | 0.02 | 30 | -1.76 | 0.827 |
| A | SALT | R | SALT | 0.12 | 0.02 | 30 | 6.43 | < 0.001 |
| A | SALT | A | BACKGROUND | -0.02 | 0.02 | 12 | -0.68 | 1 |
| A | SALT | A | TEMP | 0.03 | 0.02 | 12 | 1.18 | 0.981 |
| A | SALT | A | SALT&TEMP | 0.04 | 0.02 | 12 | 1.75 | 0.818 |
| S | SALT | R | SALT | 0.16 | 0.02 | 30 | 8.19 | < 0.001 |
| S | SALT | S | BACKGROUND | -0.02 | 0.02 | 12 | -0.68 | 1 |
| S | SALT | S | TEMP | 0.03 | 0.02 | 12 | 1.18 | 0.981 |
| S | SALT | S | SALT&TEMP | 0.04 | 0.02 | 12 | 1.75 | 0.818 |
| R | SALT | R | BACKGROUND | -0.02 | 0.02 | 12 | -0.68 | 1 |
| R | SALT | R | TEMP | 0.03 | 0.02 | 12 | 1.18 | 0.981 |
| R | SALT | R | SALT&TEMP | 0.04 | 0.02 | 12 | 1.75 | 0.818 |
| A | BACKGROUND | S | BACKGROUND | -0.03 | 0.02 | 30 | -1.76 | 0.827 |
| A | BACKGROUND | R | BACKGROUND | 0.12 | 0.02 | 30 | 6.43 | < 0.001 |
| A | BACKGROUND | A | TEMP | 0.04 | 0.02 | 12 | 1.86 | 0.762 |
| A | BACKGROUND | A | SALT&TEMP | 0.05 | 0.02 | 12 | 2.43 | 0.454 |
| S | BACKGROUND | R | BACKGROUND | 0.16 | 0.02 | 30 | 8.19 | < 0.001 |
| S | BACKGROUND | S | TEMP | 0.04 | 0.02 | 12 | 1.86 | 0.762 |
| S | BACKGROUND | S | SALT&TEMP | 0.05 | 0.02 | 12 | 2.43 | 0.454 |
| R | BACKGROUND | R | TEMP | 0.04 | 0.02 | 12 | 1.86 | 0.762 |
| R | BACKGROUND | R | SALT&TEMP | 0.05 | 0.02 | 12 | 2.43 | 0.454 |
| A | TEMP | S | TEMP | -0.03 | 0.02 | 30 | -1.76 | 0.827 |
| A | TEMP | R | TEMP | 0.12 | 0.02 | 30 | 6.43 | < 0.001 |
| A | TEMP | A | SALT&TEMP | 0.01 | 0.02 | 12 | 0.56 | 1 |
| S | TEMP | R | TEMP | 0.16 | 0.02 | 30 | 8.19 | < 0.001 |
| S | TEMP | S | SALT&TEMP | 0.01 | 0.02 | 12 | 0.56 | 1 |
| R | TEMP | R | SALT&TEMP | 0.01 | 0.02 | 12 | 0.56 | 1 |
| A | SALT&TEMP | S | SALT&TEMP | -0.03 | 0.02 | 30 | -1.76 | 0.827 |
| A | SALT&TEMP | R | SALT&TEMP | 0.12 | 0.02 | 30 | 6.43 | < 0.001 |
| S | SALT&TEMP | R | SALT&TEMP | 0.16 | 0.02 | 30 | 8.19 | < 0.001 |

**Table S6.** Model output for estimating the stressor and experimental phase effect on the necromass degradation rate of incubations with whole cells.

| Parameter | Sum Sq | Mean Sq | NumDF | DenDF | F value | P-value |
|---|---|---|---|---|---|---|
| Treatment during stressor phase | 0.06 | 0.02 | 3 | 42 | 3.1 | 0.037 |
| Phase | 0.7 | 0.35 | 2 | 42 | 53.05 | < 0.001 |

**Table S7.** Pairwise comparison of experimental phase and stressor treatment of incubations with whole cells.

| Phase1 | Treatment during stressor phase1 | Phase2 | Treatment during stressor phase2 | Estimate | SE | Df | t-ratio | P-value |
|---|---|---|---|---|---|---|---|---|
| A | SALT | S | SALT | -0.07 | 0.03 | 30 | -2.45 | 0.409 |
| A | SALT | R | SALT | 0.21 | 0.03 | 30 | 7.44 | < 0.001 |
| A | SALT | A | BACKGROUND | -0.01 | 0.03 | 12 | -0.19 | 1 |
| A | SALT | A | TEMP | -0.05 | 0.03 | 12 | -1.6 | 0.88 |
| A | SALT | A | SALT&TEMP | -0.09 | 0.03 | 12 | -2.64 | 0.352 |
| S | SALT | R | SALT | 0.28 | 0.03 | 30 | 9.89 | < 0.001 |
| S | SALT | S | BACKGROUND | -0.01 | 0.03 | 12 | -0.19 | 1 |
| S | SALT | S | TEMP | -0.05 | 0.03 | 12 | -1.6 | 0.88 |
| S | SALT | S | SALT&TEMP | -0.09 | 0.03 | 12 | -2.64 | 0.352 |
| R | SALT | R | BACKGROUND | -0.01 | 0.03 | 12 | -0.19 | 1 |
| R | SALT | R | TEMP | -0.05 | 0.03 | 12 | -1.6 | 0.88 |
| R | SALT | R | SALT&TEMP | -0.09 | 0.03 | 12 | -2.64 | 0.352 |
| A | BACKGROUND | S | BACKGROUND | -0.07 | 0.03 | 30 | -2.45 | 0.409 |
| A | BACKGROUND | R | BACKGROUND | 0.21 | 0.03 | 30 | 7.44 | < 0.001 |
| A | BACKGROUND | A | TEMP | -0.05 | 0.03 | 12 | -1.41 | 0.94 |
| A | BACKGROUND | A | SALT&TEMP | -0.08 | 0.03 | 12 | -2.45 | 0.443 |
| S | BACKGROUND | R | BACKGROUND | 0.28 | 0.03 | 30 | 9.89 | < 0.001 |
| S | BACKGROUND | S | TEMP | -0.05 | 0.03 | 12 | -1.41 | 0.94 |
| S | BACKGROUND | S | SALT&TEMP | -0.08 | 0.03 | 12 | -2.45 | 0.443 |
| R | BACKGROUND | R | TEMP | -0.05 | 0.03 | 12 | -1.41 | 0.94 |
| R | BACKGROUND | R | SALT&TEMP | -0.08 | 0.03 | 12 | -2.45 | 0.443 |
| A | TEMP | S | TEMP | -0.07 | 0.03 | 30 | -2.45 | 0.409 |
| A | TEMP | R | TEMP | 0.21 | 0.03 | 30 | 7.44 | < 0.001 |
| A | TEMP | A | SALT&TEMP | -0.03 | 0.03 | 12 | -1.04 | 0.992 |
| S | TEMP | R | TEMP | 0.28 | 0.03 | 30 | 9.89 | < 0.001 |
| S | TEMP | S | SALT&TEMP | -0.03 | 0.03 | 12 | -1.04 | 0.992 |
| R | TEMP | R | SALT&TEMP | -0.03 | 0.03 | 12 | -1.04 | 0.992 |
| A | SALT&TEMP | S | SALT&TEMP | -0.07 | 0.03 | 30 | -2.45 | 0.409 |
| A | SALT&TEMP | R | SALT&TEMP | 0.21 | 0.03 | 30 | 7.44 | < 0.001 |
| S | SALT&TEMP | R | SALT&TEMP | 0.28 | 0.03 | 30 | 9.89 | < 0.001 |

**Table S8.** Effect of experimental phase, stressor treatment, and necromass type tested with a full-factorial three-factor PERMANOVA (function adonis of R package vegan with 999 permutation). Significance codes: $p < 0.001$ ***, $p < 0.01$ **; $p < 0.05$ *.

| Factors | Df | SumsOfSqs | $R^2$ | *F*-value | *P*-value | signifi-cance |
|---|---|---|---|---|---|---|
| phase | 2 | 3.7788 | 0.25 | 37.5113 | 0.001 | *** |
| necromass | 2 | 1.9835 | 0.13 | 19.6896 | 0.001 | *** |
| stressor | 3 | 0.2748 | 0.02 | 1.8187 | 0.023 | * |
| phase*necromass | 4 | 2.0589 | 0.14 | 10.2189 | 0.001 | *** |
| phase*stressor | 6 | 0.5659 | 0.04 | 1.8725 | 0.003 | ** |
| necromass*stressor | 6 | 0.3200 | 0.02 | 1.0590 | 0.344 | |
| Phase*necromass*stressor | 12 | 0.6105 | 0.04 | 1.0101 | 0.441 | |
| residual | 104 | 5.2384 | 0.35 | | | |
| total | 139 | 14.8307 | 1.00 | | | |
